# Cross-Species Morphology Learning Enables Nucleic Acid-Independent Detection of Live Mutant Blood Cells

**DOI:** 10.1101/2025.10.20.682949

**Authors:** Sarmad Ahmad Khan, Dominik Faerber, Danielle Kirkey, Derek Stirewalt, Simon Raffel, Brandon Hadland, Michael Deininger, Florian Buettner, Helong Gary Zhao

## Abstract

In both neonates and adults, the presence of malignancy-associated mutations in peripheral blood (PB) correlates with an elevated risk of future neoplastic transformation, with certain mutations, such as *KMT2A* rearrangements, exhibiting near-complete penetrance. If feasible, pre-malignant screening could enable early intervention and even disease prevention. However, nucleic acid sequencing- and hybridization-based mutation detection have limited cost-efficiency, constraining their use in screening. Here, we introduce a computer vision platform that can identify mutant cells in fresh PB samples that carry *KMT2A-MLLT3* (a frequent mutation in pediatric and adult leukemias and detectable in newborn blood samples) or *JAK2*^*V617F*^ (a frequent mutation in myeloproliferative neoplasms and clonal hematopoiesis). This is achieved by high-throughput single-cell imaging and mutation detection by machine learning (ML)-powered morphology recognition. The ML models were developed by cross-species learning of conserved features between mutant cells from mouse genetic models and from human samples, enabling a cost-effective approach for detecting mutations in live blood cells. This platform holds promise for pre-malignant screening in asymptomatic neonates and adults with *KMT2A-MLLT3* or *JAK2*^*V617F*^mutation and is potentially generalizable to the detection other malignancy-associate mutations. Our platform provides a novel single-cell morphological data modality that complements existing single-cell genomics.

## Main Text

Mutation detection is fundamental to the molecular diagnostics of hematologic malignancies. In asymptomatic adults, the presence of malignancy-associated mutations, termed clonal hematopoiesis, is associated with increased risks of hematologic malignancies and other chronic diseases, such as cardiovascular diseases^1^. Similarly, oncogenic mutations can be detected in peripheral blood (PB) samples of newborns who later develop pediatric acute leukemia^2^. Notably, the presence of *KMT2A-* rearrangment mutations in cord blood is associated with near 100% progression to pediatric acute leukemia^2^. These underscore the potential impact of identifying early mutant lesions months to years (pediatric) or even decades (adults) before the onset of hematologic malignancies, enabling earlier intervention. Therefore, cost-efficient and high-sensitivity detection of rare mutant cells is of clinical significance.

Currently, oncogenic mutations are detected through nucleic acid-based techniques, such as DNA sequencing and *in situ* hybridization. In hematologic malignancies, next-generation sequencing (NGS) is the standard-of-practice for high-sensitivity detection of recurrent mutations. However, the high cost and technical limitations of NGS^3^ restrict its use in screening.

Machine learning (ML)-based image recognition has been shown to match or even out-perform the accuracies of medical professionals in detecting cancerous lesions, potentially assisting in clinical diagnosis^4^. In hematopathology, ML models trained on annotated blood smears have shown high accuracy in lineage classification^5^ and drug response prediction in acute myeloid leukemia (AML)^6^. With the goal of enabling pre-malignant screening, we sought to detect mutant blood cells by their native morphology, through high-throughput single-cell imaging and supervised ML. Given the impracticality of acquiring mutant cell images from patients’ PB samples (i.e. the challenge of obtaining ground-truth mutation status without compromising cellular morphology), we established a cross-species learning platform, in which mutation-specific morphological features were learned from orthologous genetic mouse models and “humanized” using one pair of mutant and normal human samples (Fig. 1A). This platform consists of two main modules: (1) mouse genetic and PDX models, and (2) single-cell imaging coupled with ML-based image recognition (Fig. 1A). Module 1 provides the training and testing materials for ML model development. In each genetic model, a single mutation, together with GFP marker, was introduced through bone marrow transplant (BMT) to generate chimeric hematopoiesis containing both mutant (GFP^+^) and normal cells (GFP^-^). Cells with GFP only were included as mock controls in testing to ensure specificity. Module 2 captures the native morphologies of fresh PB cells in three channels (brightfield, nucleus, and side scatter) and learns mutation-associated features. For better interpretability, we employed an explainable AI (XAI) approach of feature extraction followed by gradient-boosted decision tree learning (XGBoost^7^). ML models were initially developed using mouse datasets and then optimized by incorporating one pair of mutant and normal human samples in training. To ensure the robustness of our ML models, we included human samples collected from three different hospitals in external testing (Fig. 1B). Moreover, to eliminate “batch effects”, images were recorded on four different ImageStream^®^ cytometers with variable hardware configurations at four different locations (Fig. 1B). Here, we showcase our cross-species learning platform with two representative mutations relevant in both malignant and pre-malignant settings: *KMT2A-MLLT3* (also known as *MLL-*AF9, a frequent mutation in pediatric and adult leukemias^8^ and detectable in newborn PB^9^) and *JAK2*^*V617F*^ (ortholog of *Jak2*^*V617F*^, a frequent mutation in myeloproliferative neoplasms^10^ and clonal hematopoiesis^11^).

**Figure 1.**
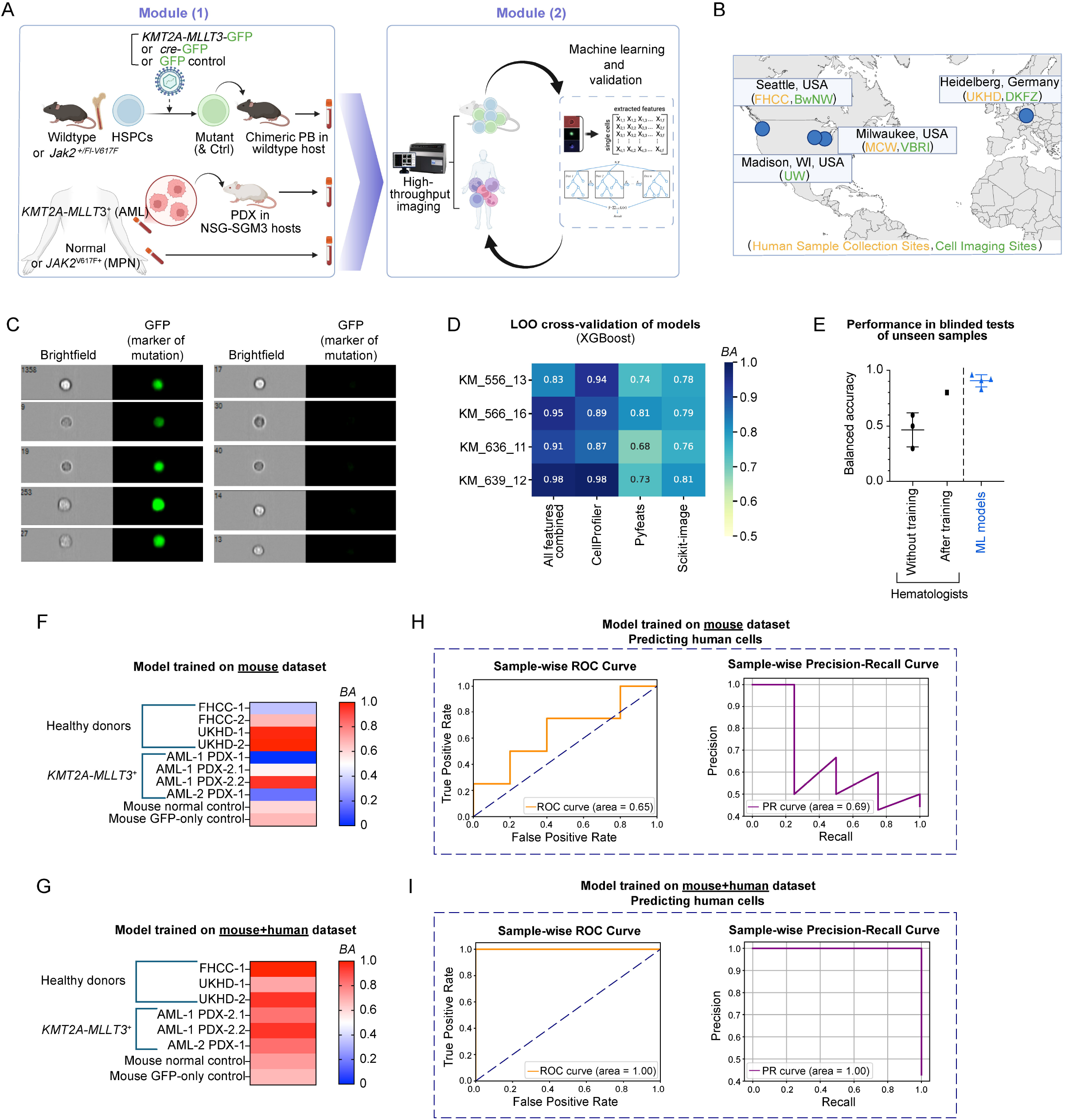
Study design and performance of ML models in detecting PB cells with *KMT2A-MLLT3* mutation. **(A)** Schematic illustration of the cross-species learning platform and the study design. CNN: convolutional neural network, MPN: myeloproliferative neoplasm. NSG: severely immunodeficient mouse strain that allows engraftment of human myeloid cells. **(B)** Study sites for human sample collection and cell imaging. FHCC: Fred Hutchinson Cancer Center, BwNW: Bloodworks Northwest, UW: University of Wisconsin-Madison, MCW: Medical College of Wisconsin, VBRI: Versiti Blood Research Institute, UKHD: Heidelberg University Hospital, DKFZ: German Cancer Research Center. **(C)** Representative images of chimeric PB samples from wildtype mouse with *KMT2A-MLLT3* mutation. Mutant cells are marked by GFP expression. Images were recorded on an ImageStream^®^ device. **(D)** Performance of XGBoost models with different combinations of cell features and different mouse datasets with *KMT2A-MLLT3* mutation. **(E)** Comparison between trained ML models and hematologists in recognizing mutant cells by native morphology. **(F)** BA of predicting mutation status in unseen human samples and controls by XGBoost models trained on mouse-only datasets with *KMT2A-MLLT3* mutation. **(G)** BA of predicting mutation status in unseen human samples and controls by XGBoost models trained on mouse datasets with *KMT2A-MLLT3* mutation plus one normal human donor sample and one *KMT2A-MLLT3*^+^ AML sample. Note that one pair of random samples (FHCC-2 and AML-1 PDX-1) from Fig. 2A were used in training. **(H, I)** Sample-wise ROC curve and PR curve of different models when applied to predicting human cells in AML PDX samples. The sample-wise median values of each parameter were used in plotting.

*KMT2A-MLLT3*^+^ leukemic cells typically exhibit myelocytic-to-monocytic phenotypes. In our mouse model, however, *KMT2A-MLLT3*^+^ PB cells showed morphologies with a variety of sizes and opacities (Fig. 1C). Despite such heterogeneity, feature-aware decision tree learning (XGBoost) achieved appreciable accuracies in mutant cell detection, with the best-performing feature extraction tool being CellProfiler^12^ (Fig. 1D, Suppl. Fig. 1C, 2A). Notably, our XAI approach outperformed widely used deep learning classifiers, including ConvNeXt and BEiT (Suppl. Fig. 1D). In comparison, three anonymous hematologists who had not been trained on unstained cell images were not able to distinguish between *KMT2A-MLLT3*^+^ and normal cells (Fig. 1E). One anonymous hematology professional was voluntarily trained on annotated images and showed improved balanced accuracy (*BA*) in a blinded test (Fig. 1E). These findings indicate that *KMT2A-MLLT3* introduces subtle alterations in native cellular morphology that can be captured by ML and are at least partially recognizable by trained human.

Models trained on <u>mouse</u> datasets exhibited inconsistent performance in distinguishing <u>human</u> *KMT2A-MLLT3*^+^ cells vs. normal counterparts (Fig. 1F), suggesting the presence species-specific morphological features. To overcome this, one pair of randomly selected *KMT2A-MLLT3*^+^ human sample (from PDX) and normal human donor sample were incorporated in training. This hybrid approach markedly improved model performance in identifying mutant human cells in unseen samples (Fig. 1G). So far, we have assessed the performance of ML models by computing the *BA* in testing, which reflects how often a correct prediction is made on each single cell. However, for future clinical translation, our primary interest lies not in cell-level predictions, but in assessing the likelihood of a given sample carrying a mutation. Therefore, we performed additional analysis at the sample level (*i*.*e*., donor/patient level), assigning each donor/patient a score that is calculated as the mean of confidence values across all cell-level predictions of that donor/patient. Subsequently, we evaluated performance metrics across all donors/patients (Fig. 1H-I). The drastically improved sample-wise ROC curve and PR curve of the mouse+human training method, compared to mouse only, underscored the necessity of human samples in the cross-species learning (Fig. 1I). Additionally, hybrid method also demonstrated high confidence in cell-level predictions across human samples (Fig. 2A). To mimic real-world applications, we tested the models’ performance with “spike-in samples” composed of mixtures of healthy donor cells and *KMT2A-MLLT3*^+^ PDX cells at pre-defined ratios. The models’ predictions showed a strong linear correlation with the actual proportions of mutant cells (Fig. 2B, Suppl. Fig. 3), indicating robust quantitative fidelity. Such linearity enables further statistical optimization of the models for large-scale retrospective clinical studies. These findings demonstrate that, in the example of *KMT2A-MLLT3*, supplementing mouse datasets with a single pair of mutant and normal human samples is sufficient to ensure high accuracy in analyzing unseen human samples.

**Figure 2.**
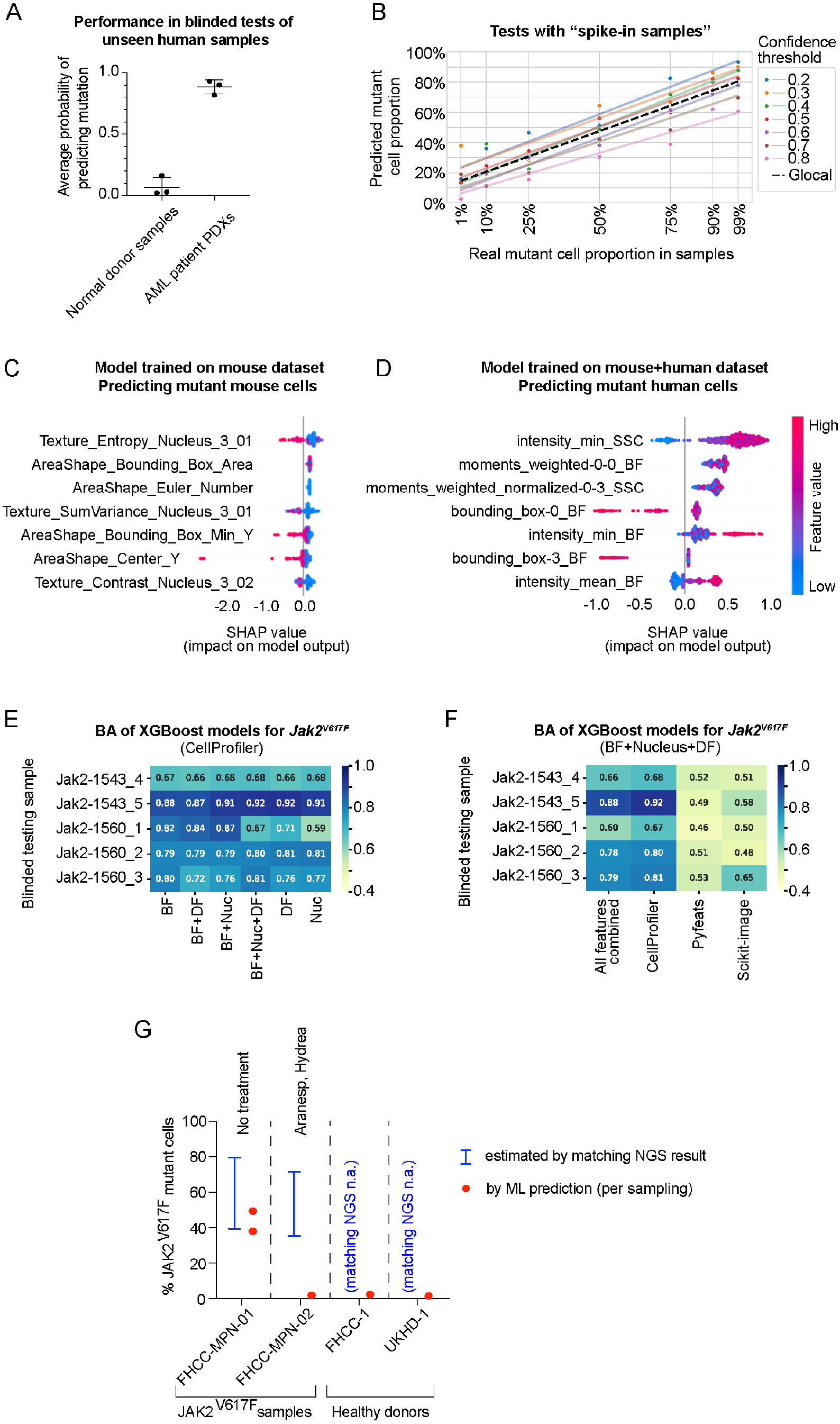
ML model’s performance with human samples, working mechanism with feature analysis, and application of the platform in detecting *Jak2*^*V617F*^/*JAK2*^*V617F*^ mutation. **(A)** Average probability of assigning “mutant” label to cells in each unseen human sample, by XGBoost models trained on mouse+human datasets with *KMT2A-MLLT3* mutation. **(B)** Performance in analyzing unseen human samples with preset proportions of mutant cells spiked in. XGBoost models were trained on mouse+human datasets with *KMT2A-MLLT3* mutation. Upper panel: XGBoost models used different confidence thresholds in calling a cell “mutant”. Lower panel: XGBoost models used different random seeds during training and LOO cross-validation. **(C)** Top features with impact (SHAP value) on the output of XGBoost models trained on mouse-only datasets with *KMT2A-MLLT3* mutation. Features were extracted from merged three-channel cell images by CellProfiler. **(D)** Top features with impact (SHAP value) on the output of XGBoost models trained on mouse+human datasets with *KMT2A-MLLT3* mutation. Features were extracted from individual channels of cell images by scikit-image. BF: brightfield, SCC: side scatter, also known as darkfield /DF. **(E, F)** Performance comparison among XGBoost models that use different channel combinations and different feature extraction methods, when tested on different *Jak2*^*V617F*^ mouse datasets during LOO cross-validation. **(G)** Test of XGBoost models (trained on mouse and healthy human samples) in predicting mutant frequency in unseen human *JAK2*^*V617F*^ patient PB samples. Estimated % of mutant cells are determined by NGS results, assuming euploidy and absence of copy number variations. Patient treatment status at the time of sample collection is labelled on top of data points. n.a.: not available.

To elucidate the specific morphological alterations induced by *KMT2A-MLLT3*, we used SHAP (SHapley Additive exPlanations) values to rank the features by their influence in mutant cell prediction. For mutant mouse cells, the top features include altered nucleus texture and increased heterogeneity in cell size (*e*.*g*. bounding box minimum) (Fig. 2C, Suppl. Fig. 4A). These are consistent with studies showing that *KMT2A-MLLT3* and AML are associated with upregulated anabolic activity, increased nuclear dynamics, and upregulation of genes controlling cell size^13-16^. Among the features shared between mouse and human cells, several stood out: increased granularity/membrane ruffles (*e*.*g*. SSC intensity and weighted moments), altered cellular texture (*e*.*g*. BF weighted moments), and opacity (*e*.*g*. BF intensity) (Fig. 2D, Suppl. Fig. 4B). These features align with the characteristic intracellular granules^17^ and monocytic morphology^18^ of *KMT2A-MLLT3*^+^ AML. Furthermore, previous reports showed that *KMT2A-MLLT3* increases both cell size and granularity in the myeloid cell line U937^19^, further validating our observations. Taken together, these results reveal substantial overlap in *KMT2A-MLLT3*-associated morphological features between mouse and human cells, supporting effective cross-species learning. Notably, since all cell types were included in ML, the identified mutation-associated features were cell type-agnostic and contribute to ML predictions in a unified, lineage-independent manner.

To explore the generalization potential of our platform, we further applied the approach to *JAK2*^*V617F*^ /*Jak2*^*V617F*^ mutation. Models trained on *Jak2*^*V617F*^ mutant and normal mouse cells yielded appreciable *BA* (Fig. 2E-F) and cell-wise ROC-AUC (Suppl. Fig. 5). To extend the model to human samples, we incorporated one normal human PB sample in training, without any mutant human samples as *JAK2*^*V617F*^ PDXs were unavailable. In blinded testing with PB samples from *JAK2*^*V617F*^ myelofibrosis and MDS/MPN patients, model predicted frequencies of *JAK2*^*V617F*^mutant cells matched the diagnostic NGS-estimated frequency for an untreated patient (FHCC-MPN-01) but disagreed with NGS-estimated frequency for a patient under antimetabolite drug treatment (FHCC-MPN-02, Fig. 2G, Suppl. Table 1). We postulate that antimetabolite drug treatment may profoundly alter the metabolic state and thus morphology of *JAK2*^*V617F*^ cells, leading to the discrepancies between model prediction and NGS estimates in FHCC-MPN-02. Whereas, due to the lack of ground truth of mutation status of each cell in human PB samples, model performance metrics were not determined (Suppl. Table 1). In future studies, we propose to use CRISPR-edited *JAK2*^*V617F*^ human HSPC xenografts in addition to mouse models and normal human samples to further improve the model’s performance.

In summary, we developed a cross-species learning platform capable of detecting mutant human cells in fresh PB samples (Suppl. Table 2). We demonstrated that two myeloid malignancy-associated mutations (*KMT2A-MLLT3* and *Jak2*^*V617F*^) induce morphological changes in mouse PB cells that are detectable by ML, independent of nucleic acids. Morphological features of mutant cells reflect the biological impact of the oncogenic mutations, such as elevated anabolic processes. Importantly, the inclusion of even a single mutant human sample during training significantly enhanced model accuracy in analyzing unseen human specimens. Our ongoing efforts aim to extend this approach to additional somatic mutations for applications in early diagnosis and pre-malignant monitoring.

## Study methods are described in the supplementary information

## Acknowledgements

We thank the members of the Hadland lab and the Meshinchi lab at FHCC, the Raffel lab at UKHD, the Deininger lab at VBRI for experimental assistance. We thank the members of the Buettner lab and the Schulz lab at DKFZ /Goethe University Frankfurt for helpful discussions. We thank the core facilities at UW-Madison, VBRI, DKFZ, and BwNW for instrumental support, and thank Tim Monahan in the FH1690 FHCC/UW Leukemia Repository Team at FHCC for coordinating patient sample collection. This work was supported by a Helmholtz International Researcher Fellowship, Fred Hutch IIRC Innovation Award, Cancer Consortium Bench-to-Bedside Adult Oncology Award (sponsored by Bezos Family Foundation) to HGZ, and NIH/NHLBI R01 grant (HL168110) to BH. This work was co-funded by the European Union (ERC, TAIPO, 101088594 to FB). Views and opinions expressed are however those of the authors only and do not necessarily reflect those of the European Union or the European Research Council. Neither the European Union nor the granting authority can be held responsible for them. SR’s research is funded by German Research Foundation (DFG) grant RA 3611/1-1.

## Authorship Contributions

HGZ and FB conceptualized the study in its current form. SK, DF, and HGZ performed ML works. HGZ and DF performed experiments, collected data, and analyzed data. MD and BH provided facilities and materials for mouse experiments. DK, DS, SR, and HGZ provided human samples. SK, HGZ, and FB wrote the manuscript with inputs from co-authors. FB, HGZ, BH, and MD provided research funding. HGZ and FB jointly supervised the study.

## References

1. Beeler, J.S., Bick, A.G., and Bolton, K.L. (2023). Genetic causes and cardiovascular consequences of clonal hematopoiesis in the UK Biobank. Nat Cardiovasc Res 2, 13–15. 10.1038/s44161-022-00198-3.

2. Marcotte, E.L., Spector, L.G., Mendes-de-Almeida, D.P., and Nelson, H.H. (2021). The Prenatal Origin of Childhood Leukemia: Potential Applications for Epidemiology and Newborn Screening. Front Pediatr 9, 639479. 10.3389/fped.2021.639479.

3. Bacher, U., Shumilov, E., Flach, J., Porret, N., Joncourt, R., Wiedemann, G., Fiedler, M., Novak, U., Amstutz, U., and Pabst, T. (2018). Challenges in the introduction of next-generation sequencing (NGS) for diagnostics of myeloid malignancies into clinical routine use. Blood Cancer J 8, 113. 10.1038/s41408-018-0148-6.

4. Tran, K.A., Kondrashova, O., Bradley, A., Williams, E.D., Pearson, J.V., and Waddell, N. (2021). Deep learning in cancer diagnosis, prognosis and treatment selection. Genome Med 13, 152. 10.1186/s13073-021-00968-x.

5. Matek, C., Krappe, S., Munzenmayer, C., Haferlach, T., and Marr, C. (2021). Highly accurate differentiation of bone marrow cell morphologies using deep neural networks on a large image data set. Blood 138, 1917–1927. 10.1182/blood.2020010568.

6. Heinemann, T., Kornauth, C., Severin, Y., Vladimer, G.I., Pemovska, T., Hadzijusufovic, E., Agis, H., Krauth, M.T., Sperr, W.R., Valent, P., et al. (2022). Deep Morphology Learning Enhances Ex Vivo Drug Profiling-Based Precision Medicine. Blood Cancer Discov 3, 502–515. 10.1158/2643-3230.BCD-21-0219.

7. Chen, T., and Guestrin, C. (2016). XGBoost: A Scalable Tree Boosting System. (ACM), pp. 785–794.

8. Meyer, C., Larghero, P., Almeida Lopes, B., Burmeister, T., Groger, D., Sutton, R., Venn, N.C., Cazzaniga, G., Corral Abascal, L., Tsaur, G., et al. (2023). The KMT2A recombinome of acute leukemias in 2023. Leukemia 37, 988–1005. 10.1038/s41375-023-01877-1.

9. Ford, A.M., Ridge, S.A., Cabrera, M.E., Mahmoud, H., Steel, C.M., Chan, L.C., and Greaves, M. (1993). In utero rearrangements in the trithorax-related oncogene in infant leukaemias. Nature 363, 358–360. 10.1038/363358a0.

10. Levine, R.L., Loriaux, M., Huntly, B.J., Loh, M.L., Beran, M., Stoffregen, E., Berger, R., Clark, J.J., Willis, S.G., Nguyen, K.T., et al. (2005). The JAK2V617F activating mutation occurs in chronic myelomonocytic leukemia and acute myeloid leukemia, but not in acute lymphoblastic leukemia or chronic lymphocytic leukemia. Blood 106, 3377–3379. 10.1182/blood-2005-05-1898.

11. Jaiswal, S., Fontanillas, P., Flannick, J., Manning, A., Grauman, P.V., Mar, B.G., Lindsley, R.C., Mermel, C.H., Burtt, N., Chavez, A., et al. (2014). Age-related clonal hematopoiesis associated with adverse outcomes. N Engl J Med 371, 2488–2498. 10.1056/NEJMoa1408617.

12. Stirling, D.R., Swain-Bowden, M.J., Lucas, A.M., Carpenter, A.E., Cimini, B.A., and Goodman, A. (2021). CellProfiler 4: improvements in speed, utility and usability. BMC Bioinformatics 22, 433. 10.1186/s12859-021-04344-9.

13. Zhu, Y., Murtadha, M., Liu, M., Caserta, E., Napolitano, O., Nguyen, L.X.T., Wang, H., Moloudizargari, M., Nigam, L., Tandoh, T., et al. (2025). Identification of CD84 as a potent survival factor in acute myeloid leukemia. J Clin Invest 135. 10.1172/JCI176818.

14. Patnana, P.K., Liu, L., Frank, D., Nimmagadda, S.C., Behrens, M., Ahmed, H., Xie, X., Liebmann, M., Wei, L., Gerdemann, A., et al. (2023). Dose-dependent expression of GFI1 alters metabolism in the haematopoietic progenitors and MLL::AF9-induced leukaemic cells. Br J Haematol 202, 1033–1048. 10.1111/bjh.18939.

15. Ni, F., Yu, W.M., Li, Z., Graham, D.K., Jin, L., Kang, S., Rossi, M.R., Li, S., Broxmeyer, H.E., and Qu, C.K. (2019). Critical role of ASCT2-mediated amino acid metabolism in promoting leukaemia development and progression. Nat Metab 1, 390–403. 10.1038/s42255-019-0039-6.

16. Cadart, C., and Heald, R. (2022). Scaling of biosynthesis and metabolism with cell size. Mol Biol Cell 33. 10.1091/mbc.E21-12-0627.

17. Goasguen, J.E., Bennett, J.M., Bain, B.J., Vallespi, T., Brunning, R., Mufti, G.J., and International Working Group on Morphology of Myelodysplastic, S. (2009). Morphological evaluation of monocytes and their precursors. Haematologica 94, 994–997. 10.3324/haematol.2008.005421.

18. Stavropoulou, V., Kaspar, S., Brault, L., Sanders, M.A., Juge, S., Morettini, S., Tzankov, A., Iacovino, M., Lau, I.J., Milne, T.A., et al. (2016). MLL-AF9 Expression in Hematopoietic Stem Cells Drives a Highly Invasive AML Expressing EMT-Related Genes Linked to Poor Outcome. Cancer Cell 30, 43–58. 10.1016/j.ccell.2016.05.011.

19. Caslini, C., Shilatifard, A., Yang, L., and Hess, J.L. (2000). The amino terminus of the mixed lineage leukemia protein (MLL) promotes cell cycle arrest and monocytic differentiation. Proc Natl Acad Sci U S A 97, 2797–2802. 10.1073/pnas.040574897.

