## Supplementary material for "Cross-Species Morphology Learning Enables Nucleic Acid-Independent Detection of Live Mutant Blood Cells": Suppl.

#### Supplementary Information

#### Methods

##### Genetic mouse models

For generating CH-like chimeric hematopoiesis with *KMT2A-MLLT3* or GFP-only mock control, C57BL/6J mouse BM cells were lentivirally transduced with *KMT2A-MLLT3*/GFP or GFP-only construct at a multiplicity of infection (MOI) of ~0.1 and transplanted into lethally irradiated isogenic littermates. For *Jak2*$^{V617F}$ mutation, B6N.129S6(SJL)-*Jak2*$^{tm1.1Ble}$/AmlyJ mouse[1] BM cells were transduced with *cre*-GFP construct at a MOI of ~0.1 to activate the V617F mutation before isogenic transplantation. PB samples with detectable GFP$^{+}$ cells (>1%) were used for imaging. Mouse studies were approved by institutional IACUC committees.

##### Human samples

For young healthy donors, PB was collected by lancet finger prick or by phlebotomy and stored in EDTA-K tubes for < 24 hours. For patients with *JAK2*$^{V617F}$, PB was collected by phlebotomy, and stored in EDTA-K tubes for < 24 hours. For AML patients with *KMT2A-MLLT3*, diagnostic BM samples were engrafted into irradiated NSG-SGM3 mice (NOD.Cg-*Prkdc*$^{scid}$ *Il2rg*$^{tm1Wjl}$ Tg(CMV-IL3,CSF2,KITLG)1Eav/MloySzJ) as patient-derived xenografts (PDXs). PB samples from PDX-carrying mice with detectable hCD45$^{+}$ cells (>1%) were used for imaging. Blood and BM donors have consented to participation in medical research. All human samples were deidentified prior to being used for experiments in this study. Use of deidentified human samples has been approved by institutional ethics committees.

##### Single-cell imaging

Single-cell images were obtained in a high-throughput fashion using Amnis ImageStream® imaging flow cytometer (Cytek Biosciences, Fremont, CA). Before imaging, 10 μL of PB sample was transferred to a sterile Eppendorf tube, and 100 μL of PBS (with 5 μM nuclear dye – DRAQ5 and 1 μg/mL dead cell dye - 7-AAD) was added to the tube to dilute the sample. 5 minutes later, diluted samples were loaded on to the Amnis ImageStream to obtain single-cell images in five channels: brightfield, side scatter (SCC / darkfield), GFP, 7AAD, and DRAQ5 (Suppl. Fig. 1A). Recorded events were gated to remove multi-cell clusters, cells out of focus, cells without nuclei, and dead cells. GFP (mutant cell label) and 7AAD (dead cell label) channels were used in cytometer gating but not in ML. ML was limited to channels of brightfield, SCC, and DRAQ5 (nucleus). Brightfield (BF), side scatter (SSC), and nuclear images of selected cells were exported as 16-bit raw pixel value tif files or as cif files using IDEAS software (Cytek Biosciences) for ML training and testing.

##### Image preprocessing

For single-cell images (tif files), the 3 channels of each cell image (BF, SSC, nuclear) were stacked into "RGB" layers of one image before use. Segmentation of cellular images was performed using a hybrid approach combining the Cellpose algorithm[2], histogram-based thresholding, and watershed-based boundary recognition.

##### Machine learning models

A collection of machine learning models was used to determine which is the best performing approach for mutation detection in our study. For feature-aware decision tree learning, gradient boosting models (*XGBoost*[3] and *CatBoost*[4]) were trained used with indicated feature sets in each experiment. Morphological and textural features of each cell were extracted using three open-source tools, namely *Pyfeats*[5], *scikit-image*[6], and *CellProfiler*[7], and the best-performing tool balanced accuracy (*BA*) was selected for each model. For CellProfiler, cif files were used and a pipeline was implemented to process cif files and stitch single-cell images into 30 × 30 grids for systematic feature extraction. ML carried out with XGBoost (Suppl. Fig.

1B). For feature-agnostic approaches, deep learning models were developed using the *Fastai*[8] framework. Deep learning architectures included ConvNext (with GELU activation)[9], ResNet (ReLU activation)[10], and Beit-large (vision transformer–based)[11]. All models were initialized with ImageNet[12] pretrained weights and trained with "RGB" stacked tif images. To address class imbalance, data augmentation was applied to the minority class (mutant cells), generating a balanced training distribution for the deep learning models, whereas for the boosting algorithms, the parameters model training parameters like scale_pos_weight in *XGBoost* and class_weights in *CatBoost* were optimized to handle the class imbalance. Model performance was evaluated by *BA* in "Leave-One-Out" (LOO) cross-validations (one biological sample insulated from training is used in blinded external validation), area under the curve of receiver operating characteristic (AUC-ROC), area under the curve of precision recall (AUC-PR). The complete code of the ML analysis is deposited on our GitHub (https://github.com/MLO-lab/leukemia-morphology-analysis). Additional description of feature extraction and ML is provided in the supplementary information.

#### ML experimental setup

Model training and testing were conducted using a leave-one-out cross-validation scheme at the individual level. For each round of training and testing, cell images of one individual mouse (with mutant and normal cells) were set aside and insulated from training and validation, which was later for external testing post-training. The remaining cell images from the other mice were split at 0.9/0.1 and used for model training and validation. We compared the balanced accuracies of all deep learning models trained with images and decision tree learning models trained with different extracted features sets to determine the best performing ML model for each classification task in our study. In all tasks, *XGBoost* model had the highest balanced accuracies across different datasets, and was used for cross-species learning in the bulk of our study. Despite the high performance of *XGBoost* model trained on mouse only image datasets in tasks of classifying unseen mouse cells, it showed uneven performances in tasks of classifying human cell images. To improve the performance of cross-species learning, we included one pair of mutant and normal human samples (for just one normal human sample for *JAK2*$^{V617F}$) into the mouse image datasets as the training material. This latter approach significantly improved the accuracy of trained *XGBoost* models in classifying unseen human cells, and was assigned as the standard approach of our cross-species learning platform.

#### Reference


1. Mullally, A., Lane, S.W., Ball, B., Megerdichian, C., Okabe, R., Al-Shahrour, F., Paktinat, M., Haydu, J.E., Housman, E., Lord, A.M., et al. (2010). Physiological Jak2V617F expression causes a lethal myeloproliferative neoplasm with differential effects on hematopoietic stem and progenitor cells. Cancer Cell *17*, 584-596. 10.1016/j.ccr.2010.05.015.
2. Stringer, C., Wang, T., Michaelos, M., and Pachitariu, M. (2021). Cellpose: a generalist algorithm for cellular segmentation. Nat Methods *18*, 100-106. 10.1038/s41592-020-01018-x.
3. Chen, T., and Guestrin, C. (2016). XGBoost: A Scalable Tree Boosting System. (ACM), pp. 785–794.
4. Anna Veronika Dorogush, V.E., Andrey Gulin (2018). CatBoost: gradient boosting with categorical features support. arXiv *1810.11363*.
5. Giakoumoglou, N. (2021). PyFeats: Open-source software for image feature extraction.
6. van der Walt, S., Schonberger, J.L., Nunez-Iglesias, J., Boulogne, F., Warner, J.D., Yager, N., Gouillart, E., Yu, T., and scikit-image, c. (2014). scikit-image: image processing in Python. PeerJ *2*, e453. 10.7717/peerj.453.

7. Stirling, D.R., Swain-Bowden, M.J., Lucas, A.M., Carpenter, A.E., Cimini, B.A., and Goodman, A. (2021). CellProfiler 4: improvements in speed, utility and usability. BMC Bioinformatics *22*, 433. 10.1186/s12859-021-04344-9.
8. Jeremy Howard, S.G. (2020). fastai: A Layered API for Deep Learning. arXiv *2002.04688*.
9. Liu, Z., Mao, H.Z., Wu, C.Y., Feichtenhofer, C., Darrell, T., and Xie, S.N. (2022). A ConvNet for the 2020s. Proc Cvpr Ieee, 11966-11976. 10.1109/Cvpr52688.2022.01167.
10. He, K.M., Zhang, X.Y., Ren, S.Q., and Sun, J. (2016). Deep Residual Learning for Image Recognition. 2016 Ieee Conference on Computer Vision and Pattern Recognition (Cvpr), 770-778. 10.1109/Cvpr.2016.90.
11. Hangbo Bao, L.D., Songhao Piao, Furu Wei (2022). BEiT: BERT Pre-Training of Image Transformers. arXiv *2106.08254*.
12. Deng, J., Dong, W., Socher, R., Li, L.J., Li, K., and Li, F.F. (2009). ImageNet: A Large-Scale Hierarchical Image Database. Cvpr: 2009 Ieee Conference on Computer Vision and Pattern Recognition, Vols 1-4, 248-255. DOI 10.1109/cvpr.2009.5206848.

**Supplementary Table 1.** Patient characteristics for *JAK2*$^{V617F}$ fresh PB samples.

| **Sample ID** | **FHCC-MPN-01** | **FHCC-MPN-02** |
|---|---|---|
| **Diagnosis** | Myelofibrosis | MDS/MPN-SF3B1 |
| **Collection Date** | 2024-10-29 | 2025-03-31 |
| **Treatment** | None | Aranesp, Hydrea |
| **NGS result** | KRAS (p.G12V, NM_004985.3:c.35G>T, VAF 48%);<br>**JAK2** (p.V617F, NM_004972.3:c.1849G>T, VAF 40%);<br>ASXL1 (p.E635Rfs*15, NM_015338.5:c.1900_1922del, VAF 10%) | DNMT3A (p.P625Lfs*26, NM_022552.4:c.1874del, VAF 39%);<br>TET2 (p.L1899Sfs*9, NM_001127208.2:c.5695del, VAF 38%);<br>**JAK2** (p.V617F, NM_004972.3:c.1849G>T, VAF 36%) |
| **Estimated *JAK2*$^{V617F}$ cell frequency** | 40~80%* | 36~72%* |

**Note: *** assuming euploidy and absence of copy number variations of *JAK2* gene.

**Supplementary Table 2.** ML model specifications and performances in all experiments. (n.d.: not determined, due to lack of ground truth of single-cell mutation status)

| | ***KMT2A-MLLT3*** | | ***JAK2$^{V617F}$*** | |
|---|---|---|---|---|
| **Training datasets** | Mouse (4) | Mouse+Human (4 + 1) | Mouse (4) | Mouse+Human (4 + 1) |
| **Testing datasets** | Mouse | Human | Mouse | Human |
| **Best feature extraction tool** | CellProfiler | Scikit-image | CellProfiler | n.d. |
| **Best channel combination** | Brightfield + Nucleus + SSC | Brightfield + SSC | Brightfield + Nucleus | n.d. |
| **BA in cross-validations** | 0.83 ~ 0.96 | 0.71 ~ 0.99 | 0.68 ~ 0.91 | n.d. |
| **AUC-ROC** | Sample-wise 0.92 ~ 1.00 | Overall 0.93 | Sample-wise 0.73 ~ 0.97 | n.d. |
| **AUC-PR** | | 0.74 | | n.d. |

**Supplementary Figure Legends**

**Supplementary Figure 1.** Flow cytometry gating strategy, ML process, and representative single-cell images in raw form and ML-ready form. **(A)** Events captured on ImageStream imaging cytometer were gated sequentially to remove speedbeads, out-of-focus cells, dead cells. In the final gates, $GFP^+$ and $GFP^-$ cells were recorded for ML, whose representative images are shown below. All channels were recorded after color compensations. Mathematical representation of the ML process is shown in the lower panel. **(B)** Representative images of cells used for ML training, their labels, and their predictions are shown. Selected channels were given pseudo-colors and combined into RGB images in tiff format.

**Supplementary Figure 2.** Sample-wise ROC curve **(A)** and feature importance **(B)** for the *KMT2A-MLLT3* mouse dataset.

**Supplementary Figure 3.** Overall feature importance for the prediction of *KMT2A-MLLT3*$^+$ mouse samples **(A)** and *KMT2A-MLLT3*$^+$ human samples **(B).**

**Supplementary Figure 4.** Sample-wise ROC curve **(A)** and feature importance **(B)** for the *Jak2*$^{+/V617F}$ mouse dataset.

### Supplementary Figure 1

**A**

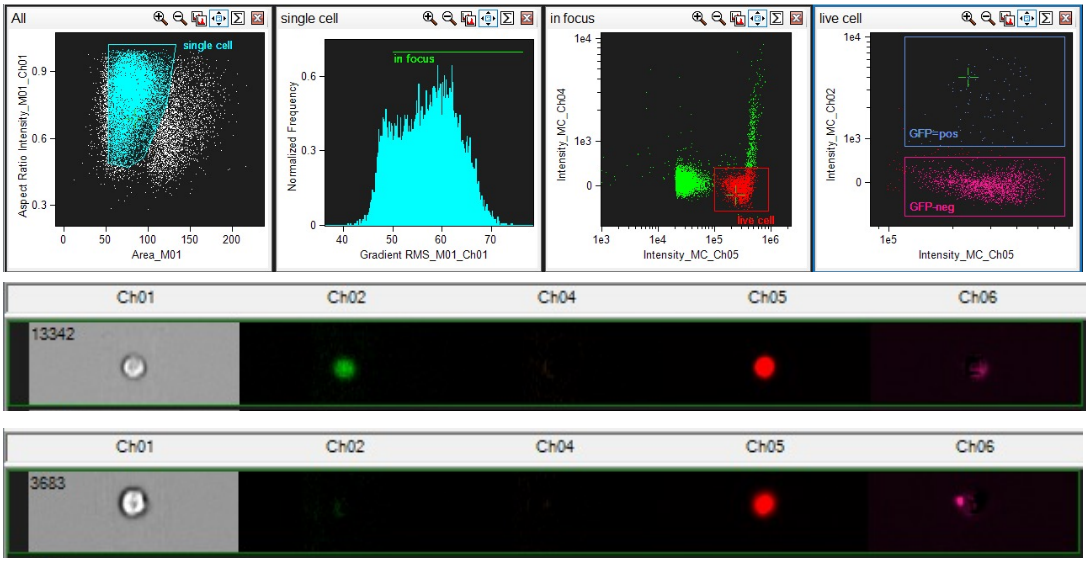


Input image: $I \in \mathbb{R}^{m \times n}$

Feature extraction (CellProfiler / scikit-image): $\varphi : \mathbb{R}^{m \times n} \to \mathbb{R}^d, \quad \mathbf{x} = \varphi(I)$

Classifier (XGBoost): $f : \mathbb{R}^d \to \{0, 1\}, \quad \hat{y} = f(\mathbf{x})$

Process: $I \xrightarrow{\varphi} \mathbf{x} \xrightarrow{f} \hat{y}$

**B**

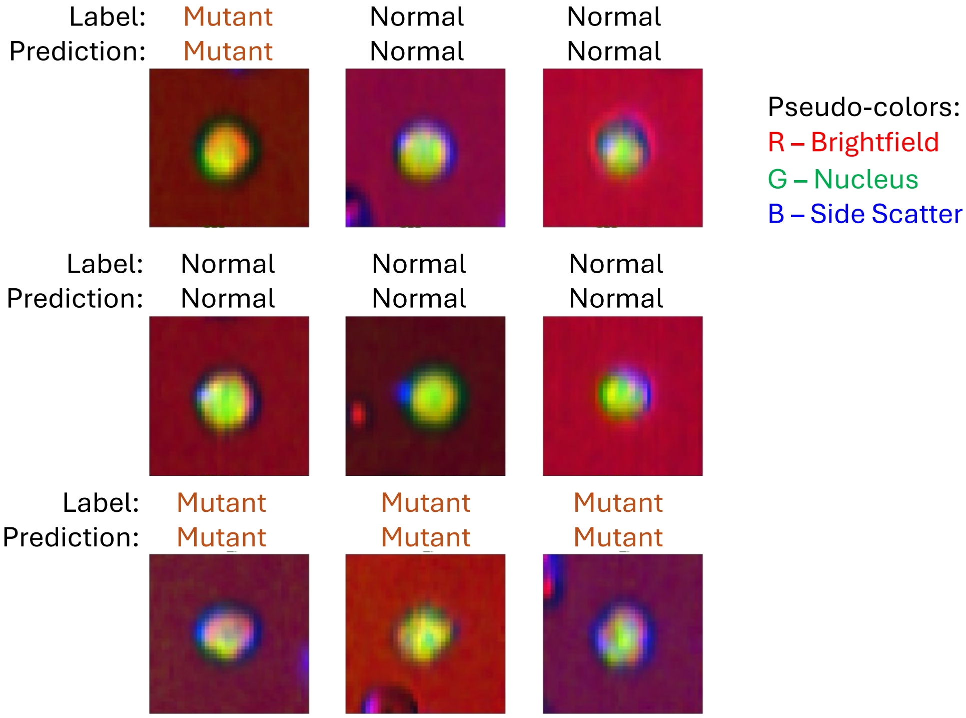

### Supplementary Figure 2

**A**

Sample-wise ROC curve
(ML models trained on mouse datasets)

*KMT2A-MLLT3*+ mouse-556 | *KMT2A-MLLT3*+ mouse-566 | *KMT2A-MLLT3*+ mouse-636

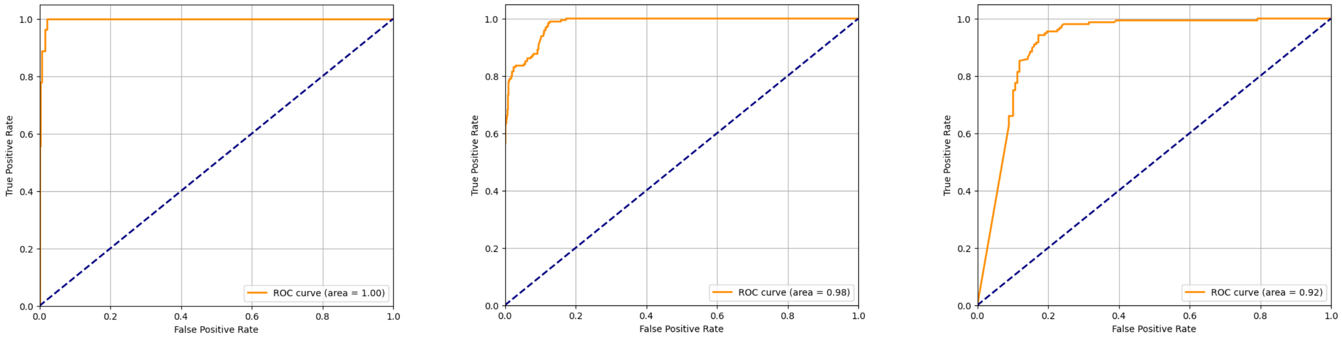


**B**

Sample-wise feature importance to model predictions
(ML models trained on mouse datasets)

*KMT2A-MLLT3*+ mouse-556 | *KMT2A-MLLT3*+ mouse-566 | *KMT2A-MLLT3*+ mouse-636

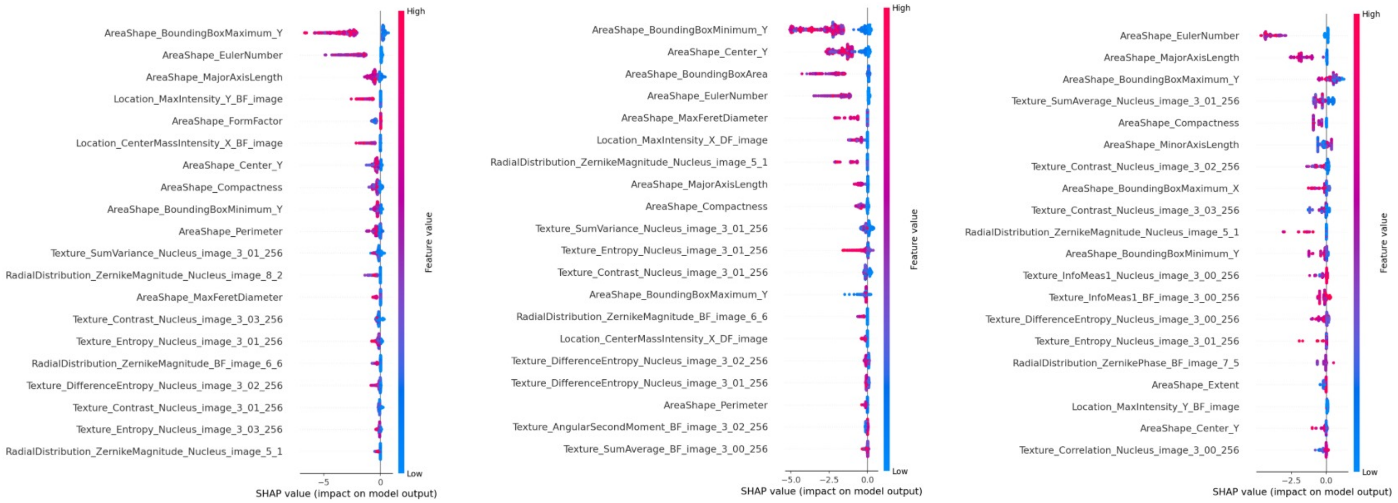

### Supplementary Figure 3

**A** Overall feature importance to prediction of unseen *KMT2A-MLLT3*+ mouse samples (ML models trained on mouse datasets)

**B** Overall feature importance to prediction of unseen *KMT2A-MLLT3*+ human samples (ML models trained on mouse+human datasets)

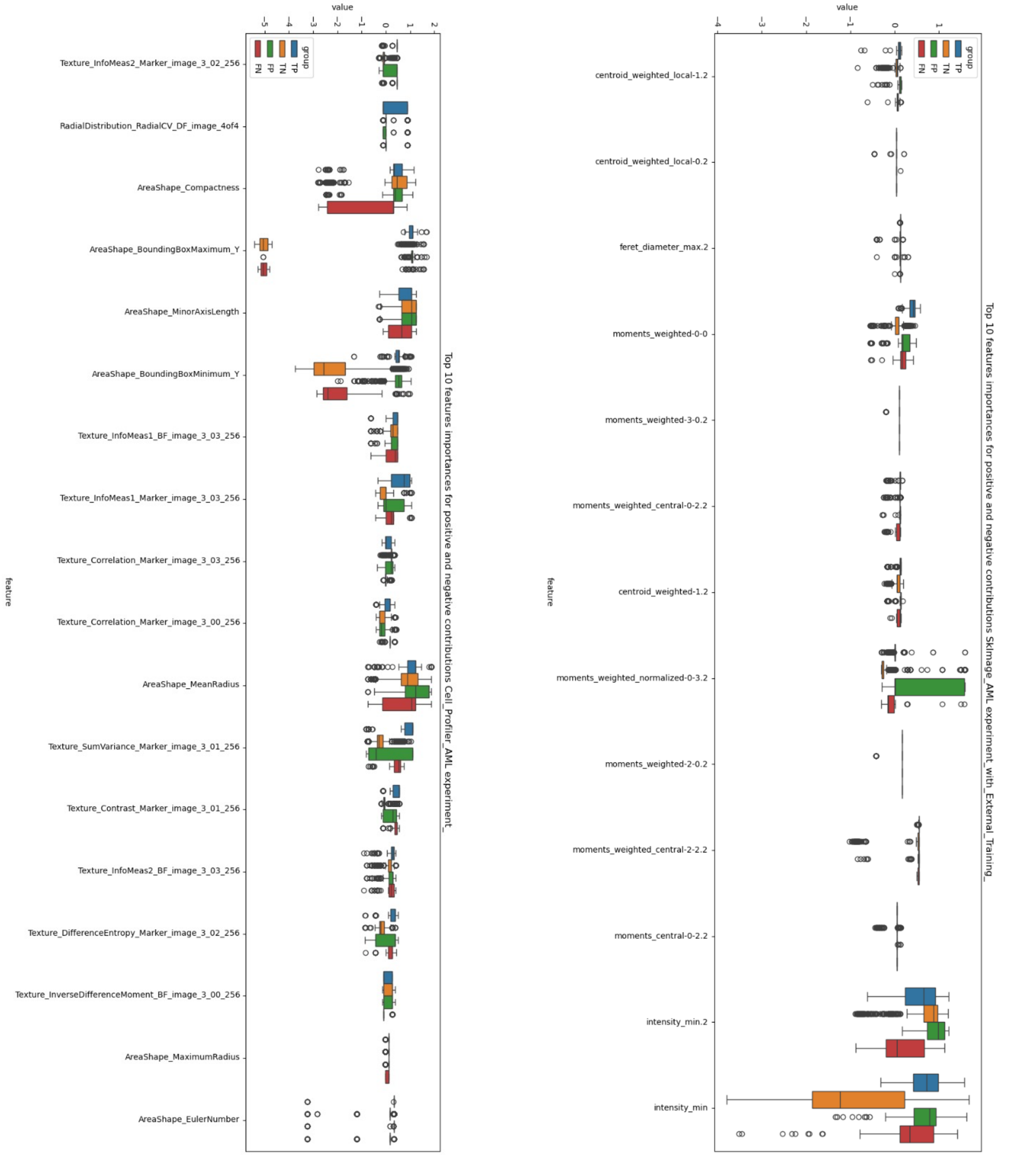

### Supplementary Figure 4

**A**

Sample-wise ROC curve
(ML models trained on mouse datasets)

*Jak2*$^{+/V617F}$ mouse-1543

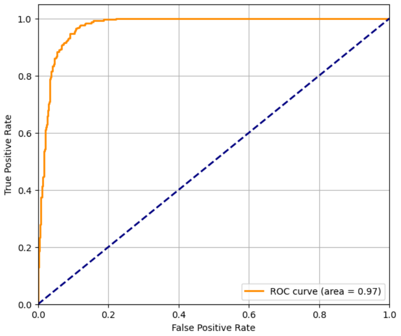


*Jak2*$^{+/V617F}$ mouse-1560

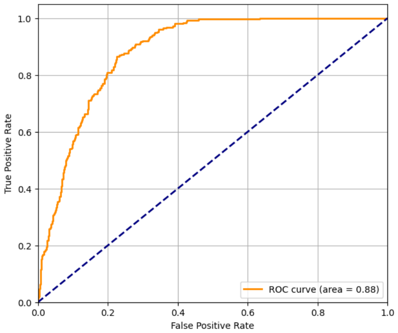


**B**

Sample-wise feature importance to model predictions
(ML models trained on mouse datasets)

*Jak2*$^{+/V617F}$ mouse-1543

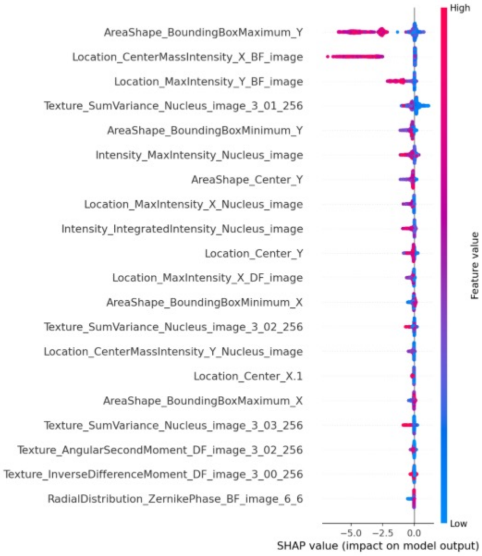


*Jak2*$^{+/V617F}$ mouse-1560

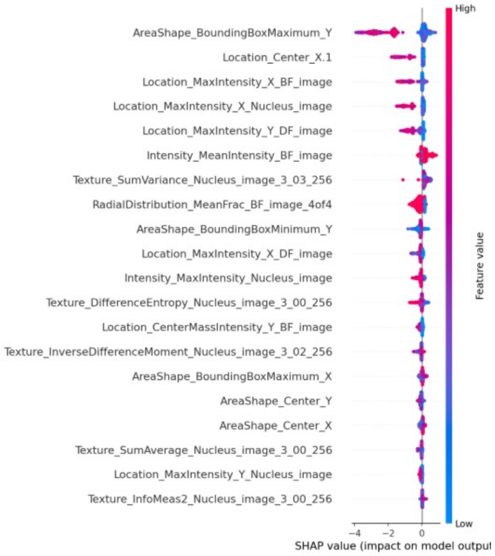
